# The coronavirus nsp14 exoribonuclease interface with the cofactor nsp10 is essential for efficient virus replication and enzymatic activity

**DOI:** 10.1101/2024.09.26.615217

**Authors:** Samantha L. Grimes, Brook E. Heaton, Mackenzie L. Anderson, Katie Burke, Laura Stevens, Xiaotao Lu, Nicholas S. Heaton, Mark R. Denison, Jordan Anderson-Daniels

## Abstract

Coronaviruses (CoVs) encode nonstructural proteins (nsps) 1-16, which assemble to form replication-transcription complexes that function in viral RNA synthesis. All CoVs encode a proofreading 3’-5’ exoribonuclease (ExoN) in nsp14 (nsp14-ExoN) that mediates proofreading and high-fidelity replication and is critical for other roles in replication and pathogenesis. The *in vitro* enzymatic activity of nsp14 ExoN is enhanced in the presence of the cofactor nsp10. We introduced alanine substitutions in nsp14 of murine hepatitis virus (MHV) at the nsp14-10 interface and recovered mutant viruses with a range of impairments in replication and *in vitro* biochemical exonuclease activity. Two of these substitutions, nsp14 K7A and D8A, had impairments intermediate between WT-MHV nsp14 and the known ExoN(-) D89A/E91A nsp14 catalytic inactivation mutant. All introduced nsp14-10 interface alanine substitutions impaired *in vitro* exonuclease activity. Passage of the K7A and D8A mutant viruses selected second-site non-synonymous mutations in nsp14 associated with improved mutant virus replication and exonuclease activity. These results confirm the essential role of the nsp14-nsp10 interaction for efficient enzymatic activity and virus replication, identify proximal and long-distance determinants of nsp14-nsp10 interaction, and support targeting the nsp14-10 interface for viral inhibition and attenuation.

**IMPORTANCE:** Coronavirus replication requires assembly of a replication transcription complex composed of nonstructural proteins (nsp), including polymerase, helicase, exonuclease, capping enzymes, and non-enzymatic cofactors. The coronavirus nsp14 exoribonuclease mediates several functions in the viral life cycle including genomic and subgenomic RNA synthesis, RNA recombination, RNA proofreading and high-fidelity replication, and native resistance to many nucleoside analogs. The nsp-14 exonuclease activity *in vitro* requires the non-enzymatic co-factor nsp10, but the determinants and importance the nsp14-10 interactions during viral replication have not been defined. Here we show that for the coronavirus murine hepatitis virus, nsp14 residues at the nsp14-10 interface are essential for efficient viral replication and *in vitro* exonuclease activity. These results shed new light on the requirements for protein interactions within the coronavirus replication transcription complex, and they may reveal novel non active-site targets for virus inhibition and attenuation.

## INTRODUCTION

Coronaviruses (CoVs) cause endemic, epidemic, and pandemic diseases in humans and animals and demonstrate frequent cross-species movement and zoonotic diseases. Thus, determining fundamental mechanisms of replication is critical to understanding their evolution and targeting for inhibition and attenuation. CoVs are members of the order *Nidovirales,* and possess the largest known and most complex single stranded (+)RNA genomes of viruses known to infect humans and animals (1,2). CoVs have significantly lower basal mutation rates than other RNA viruses; this relative higher fidelity of replication is mediated by the proofreading exoribonuclease (ExoN) which is conserved in the large nidoviruses (3,4). ExoN-mediated proofreading has been proposed to be a key factor in the ability of large nidoviruses to faithfully maintain genetic integrity of large genomes in the setting of selective pressures (3,5).

The 5’ two-thirds of the CoV (+)RNA genome encodes 16 nonstructural proteins (nsps1-16) (Fig. 1) (6). Coronavirus replication in cells is initiated by the translation of the input (+)RNA genome into two co-amino-terminal polyproteins encoding nsp1-16, followed by proteolytic maturation processing by two or three protease activities within the translated polyproteins (7–9). Structural, biochemical, and genetic studies have shown or predicted roles for multiple nsps in the formation and function of the membrane associated replication-transcription complexes (RTC) that mediate all stages of viral RNA synthesis (6). These include: nsp12 RNA-dependent RNA polymerase (nsp12-RdRp) and its cofactors nsp7 and nsp8; nsp13 helicase-ATPase involved in RNA unwinding; nsp14 exonuclease and N7-methyltransfersase activities; nsp15 endoribonuclease; and nsp16 2’-O methyltransferase (6,10–13).

**Figure 1.**
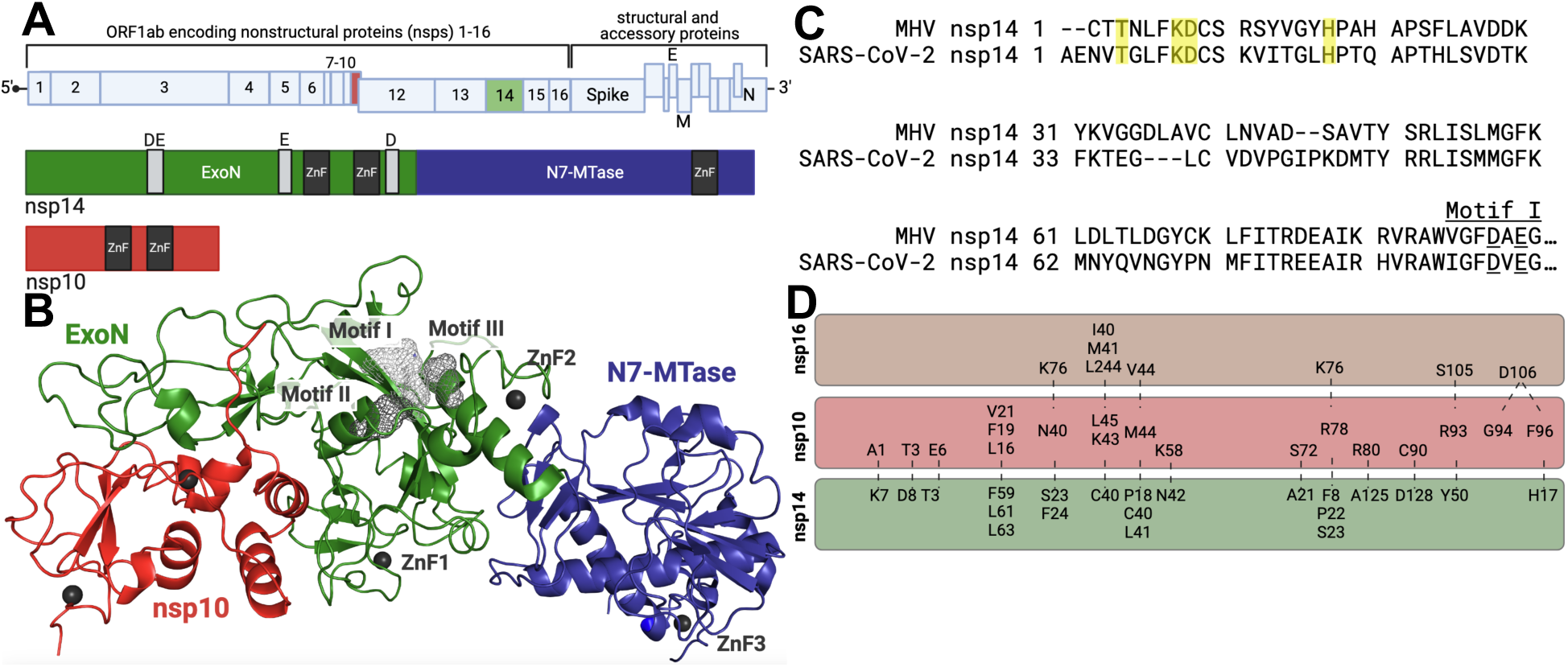
Coronavirus genome, nonstructural protein 14 (nsp14) structure, conservation, and interface with nsp10. **(A)** Schematic of the SARS-CoV-2 genome, showing open reading frame (ORF) 1ab which encodes nonstructural proteins 1-16, as well as the region of the genome encoding structural and accessory proteins. Colored are nsp10 (red) and nsp14 (green/blue). Below, a linear depiction of the SARS-CoV-2 nsp14 domains showing the 3’-5’ exoribonuclease [ExoN] (green) and N7 methyltransferase [N7-MTase] (blue) domains. The ExoN catalytic motifs for SARS-CoV-2 are shown in light gray (Motif I: D90 and E92; Motif II: E191, Motif III: D273). The zinc fingers are shown in dark gray. Shown also is nsp10, a cofactor for both nsp14 and nsp16 with the two zinc fingers shown in dark gray. **(B)** The CryoEM structure of the SARS-CoV-2 nsp14 bound to nsp10 (PDB: 7N0B). Colors are the same as in panel A, with domains and motifs labeled. **(C)** Amino acid alignment between the MHV nsp14 (YP_009924354.1) and SARS-CoV-2 (YP_009724389), with residues of interest highlighted in yellow. **(D)** Diagram of the predicted nsp10-nsp14 interface and the nsp10-nsp16 interface in MHV. Dashed lines signify likely interactions based on biochemical and structural studies.

The CoV nsp14 has two distinct functional domains, the amino-terminal 3’-to-5’ exoribonuclease (nsp14-ExoN) and the carboxy-terminal N7 methyltransferase (nsp14-N7-MTase) (Fig. 1) The nsp14-ExoN has many functions during CoV replication and pathogenesis. The exonuclease activity of ExoN is dependent on 4 conserved active residues in 3 motifs: motif I-DE, motif II-E, and motif III-DH. The nsp14 exonuclease hydrolyses ssRNA and dsRNA in in a 3’-to-5’ direction (14) and excises 3’ single-nucleotide mismatches in template RNA (15). ExoN is required for WT viral RNA accumulation and subgenomic RNA synthesis (14). Viable mutants of SARS-CoV and murine hepatitis virus (MHV) with alanine substitutions at ExoN motif I (DE) residues [hereafter ExoN(-)] have: 1) defects in virus replication and RNA synthesis (3,16); 2) up to 20-fold increased mutation frequency compared to WT parental viruses, and more consistent with higher mutation rates of other RNA viruses (3,16); 3) increased sensitivity to nucleoside analogs including 5-fluorouracil (5-FU), remdesivir, and molnupiravir (5,17,18); 4) decreased and altered recombination functions during genomic and subgenomic RNA synthesis (19); 5) increased sensitivity to interferon (20,21); 6) loss of fitness compared to WT virus (22); and 7) *in vivo* attenuation of an ExoN(-) mutant of lethal mouse adapted SARS-CoV (23). More recent studies in porcine epidemic diarrhea virus (PEDV), SARS-CoV and SARS-CoV-2 support the role of nsp14 as an innate immune antagonist (20,24,25). Finally, investigators in our lab and others have been unable to recover ExoN(-) mutants of the betacoronaviruses MERS-CoV and SARS-CoV-2 and the alpha CoV HCoV-229E, suggesting nsp14-ExoN likely serves additional essential functions in these viruses. (14,26).

Studies of nsp14 structure and exonuclease activity have demonstrated that nsp10, a small non-enzymatic protein, has specific interactions with nsp14 (27–30). The *in vitro* exonuclease activity of nsp14-ExoN requires or is enhanced by nsp10, as is stability and resolution of nsp14 structures solved by crystallography or cryo-EM (Fig. 1)) (28–31). Nsp10 also is a required cofactor for the *in vitro* activity and stable structure of nsp16 2’-O-methyltransferase (nsp16-2’-O-MTase) (32). The residues of nsp10 that interact at an interface with nsp14 share a partially overlapping footprint with the residues that interact with nsp16-2’-OMTase (Fig. 1) (29). Amino acid substitutions at nsp10 residues that interface with nsp14 limit nsp10 association with nsp14, impair nsp14-exonuclease activity *in vitro*, and may impact N7-MTase activity (33). In our genetic studies with MHV, alanine substitutions in nsp10 at some residues proposed to interface with nsp14 were non-viable, while others were viable but had altered sensitivity to nucleoside analogs under different thermal conditions (34). However, genetic and biochemical studies of mutations in nsp10 are limited for conclusions about impact of the nsp14-10 interface because of our evolving but limited understanding of the structural and biochemical interactions of nsp10, nsp14, and nsp16 during virus replication, interactions in the larger RTC during viral RNA synthesis, and impact on other nsp14 functions. A recent study reports a heterotrimeric structure of SARS-CoV-2 nsp10-14-16 (35); however, during infection, assembly of CoV RTC proteins, and genome replication, it is not known if nsp10 molecules shuttle or compete between nsp14 and nsp16, or if distinct nsp10 molecules exclusively interact with one protein or the other. Therefore, understanding the role of the nsp14-10 interface during infection requires genetic and biochemical approaches targeting nsp14.

In this study, we show that alanine substitutions in MHV nsp14 residues at the nsp14-nsp10 interface variably impact virus replication, exonuclease activity, and nucleoside analog sensitivity. Two nsp14 substitutions, K7A and D8A, had the greatest impact on replication, exonuclease activity, and fitness, but did not increase sensitivity to nucleoside analogues. Passage of these mutants partially restored virus replication and selected for second-site substitutions in nsp14 that, when reintroduced into the mutant virus background, independently partially compensated replication and biochemical activities but remained less fit than WT MHV. Together these data demonstrate that the nsp14-nsp10 interface, and specifically the nsp14 residues at this interface, are key regulators of nsp14 functions, but have functions in addition to facilitating exonuclease activity.

## RESULTS

We introduced mutations coding for alanine substitutions at four residues in the murine hepatitis virus (MHV) nsp14 at the proposed nsp14-10 interface that were conserved across divergent CoVs: T3, K7, D8, and H17 (Fig. 1). Using reverse genetics, we recovered MHV harboring the intended mutations and generated low passage stocks that were used for all experiments presented. Both T3A and H17A mutant viruses produced WT-like plaques in DBT-9 cells, while K7A and D8A mutant viruses produced small and medium-size plaques. We compared the replication kinetics of the mutant viruses to that of WT MHV and to the well-characterized, catalytically inactive nsp14-ExoN(-) mutant (D89A /E91A) (Fig. 2). T3A and H17A viruses began replication with similar kinetics as WT MHV; however, both mutant viruses had peak titers less than WT-MHV. In contrast, the K7A and D8A viruses demonstrated a 2h delay to onset of exponential replication compared to WT and had impaired peak titers 10-50 fold less than WT-MHV. All four mutant viruses demonstrated earlier onset of exponential replication and increased peak titers compared to nsp14-ExoN(-). These data indicate that nsp14 residues at the nsp10 interface contribute to efficient MHV replication.

**Figure 2.**
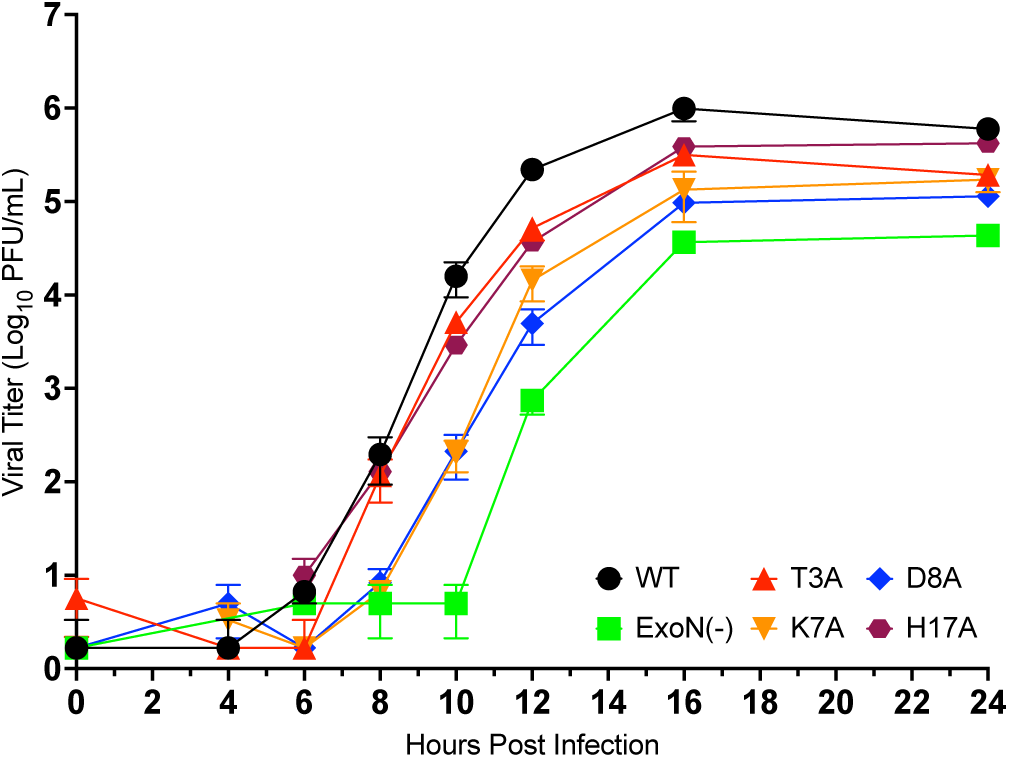
Replication of nsp14-10 interface mutants. Replication of MHV-WT, ExoN(-), nsp14-T3A, nsp14-K7A, nsp14-D8A, and nsp14-H17A. DBT-9 cells were infected with the indicated virus at an MOI of 0.01 PFU/cell. Supernatant samples were collected at the indicated times, and the virus titer was determined by plaque assay. Data shown are from three independent experiments, each with three replicates. Points represent experiment means (n=3) ± SEM.

### Competitive fitness of mutant viruses

We next tested the fitness of T3A, K7A, D8A, and H17A viruses using a coinfection competitive fitness assay (36–38) (Fig. 3). Three independent lineages of mutant and WT viruses were coinfected with competitor WT MHV encoding a genetic barcode of seven silent mutations in the nsp2 coding domain (WT-BC). The resulting viral populations were then passaged an additional three times at a constant MOI of 0.1 PFU/cell. Viral population RNA was extracted from the supernatants of each passage, and primers detecting either the barcoded (WT-BC infection) or nonbarcoded (WT control, mutant competitor) nsp2 cDNA were used in RT-qPCR reactions. The ratio of nonbarcoded to barcoded cDNA was plotted over passage number. All mutant virus cDNAs were less abundant than WT by passage 1 with varying degrees of downward trends through passage 4. We analyzed the competitor/WT-BC ratio by linear regression to determine the competitive fitness of each mutant relative to WT MHV. T3A, K7A, and D8A mutant competitors had significant reductions in fitness relative to WT, while the fitness of H17A was not significantly different from WT. These data are consistent with the replication kinetics data.

**Figure 3.**
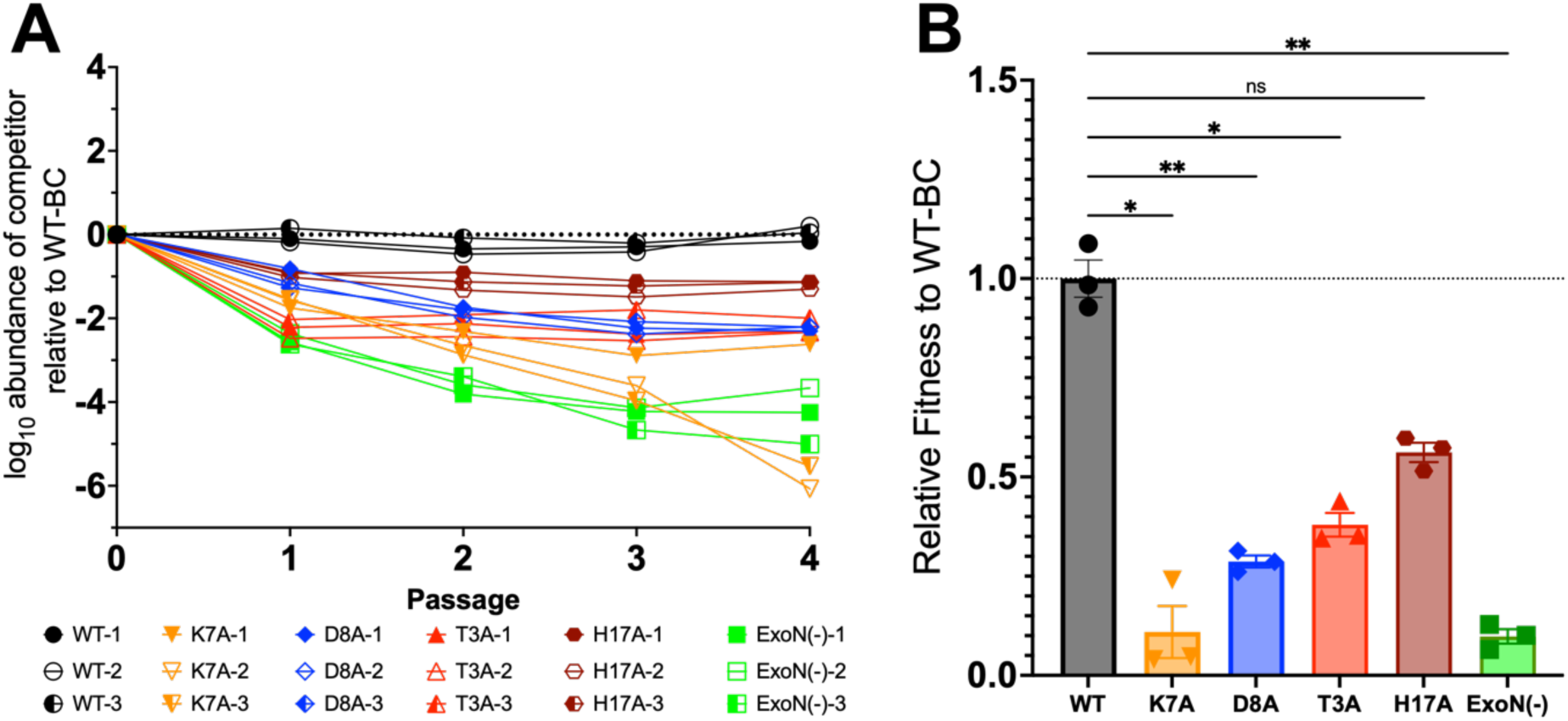
Competitive fitness of nsp14-10 interface mutants. **(A)** DBT-9 cells were co-infected with a barcoded WT MHV and a nonbarcoded WT, K7A, D8A, T3A, H17A, or ExoN(-) at a combined MOI of 0.1 PFU/cell. The resulting supernatants were passaged 4 times. The relative quantities of barcoded and nonbarcoded cDNAs were plotted over passage for the three independent lineages of each competition, normalized to the input amounts. **(B)** Linear regression from panel A was used to determine the relative fitness for each nonbarcoded virus. Individual data means are graphed (n=3) ± SEM. Statistical significance determined by one-way ANOVA with Dunnett’s multiple-comparison test. * p < 0.05; ** p < 0.01, ns not significant.

### Mutant virus sensitivity to nucleoside analogues

WT MHV has native resistance to several nucleoside analogues, a trait that is associated with intact nsp14 catalytic activity. The MHV-ExoN(-) mutant virus has increased inhibition by multiple nucleoside analogs, including 5-fluorouracil (5-FU), consistent with loss of proofreading or enhanced selectivity with loss of ExoN function (5). To determine if the nsp14-10 interface mutants share this phenotype with MHV-ExoN(-), we compared the 5-FU sensitivities of WT-MHV, MHV-ExoN(-), and the T3A, K7A, D8A, and H17A mutant viruses (Fig. 4). WT-MHV was unaffected by 5-FU concentrations up to 200μM, while MHV-ExoN(-) was inhibited in a dose-dependent manner with complete inhibition by 100μM 5-FU. In contrast, the T3A, K7A, D8A, and H17A viruses all retained WT-like resistance to 5-FU. We next tested mutant sensitivity to the nucleoside analog EIDD-1931, a cytidine analog and active form of the antiviral molnupiravir with broad spectrum activity against CoVs, and which functions as an RNA mutagen during replication. Consistent with previous experiments, WT MHV demonstrated dose-dependent inhibition by EIDD-1931, and MHV-ExoN(-) had enhanced sensitivity to EIDD-1931 at all concentrations tested up to 15μM (18). In contrast, the T3A, K7A, D8A, and H17A mutant viruses retained WT-like inhibition by EIDD-1931 up to 2µM, with increases in inhibition between 2µM and 15µM. They were 100-fold less sensitive to EIDD-1931 at 2µM compared to MHV-ExoN(-), and mutant virus infection was detectible at all EIDD-1931 concentrations tested. Thus, the interface substitutions did not directly impact the sensitivity to either 5-FU or EIDD-1931.

**Figure 4.**
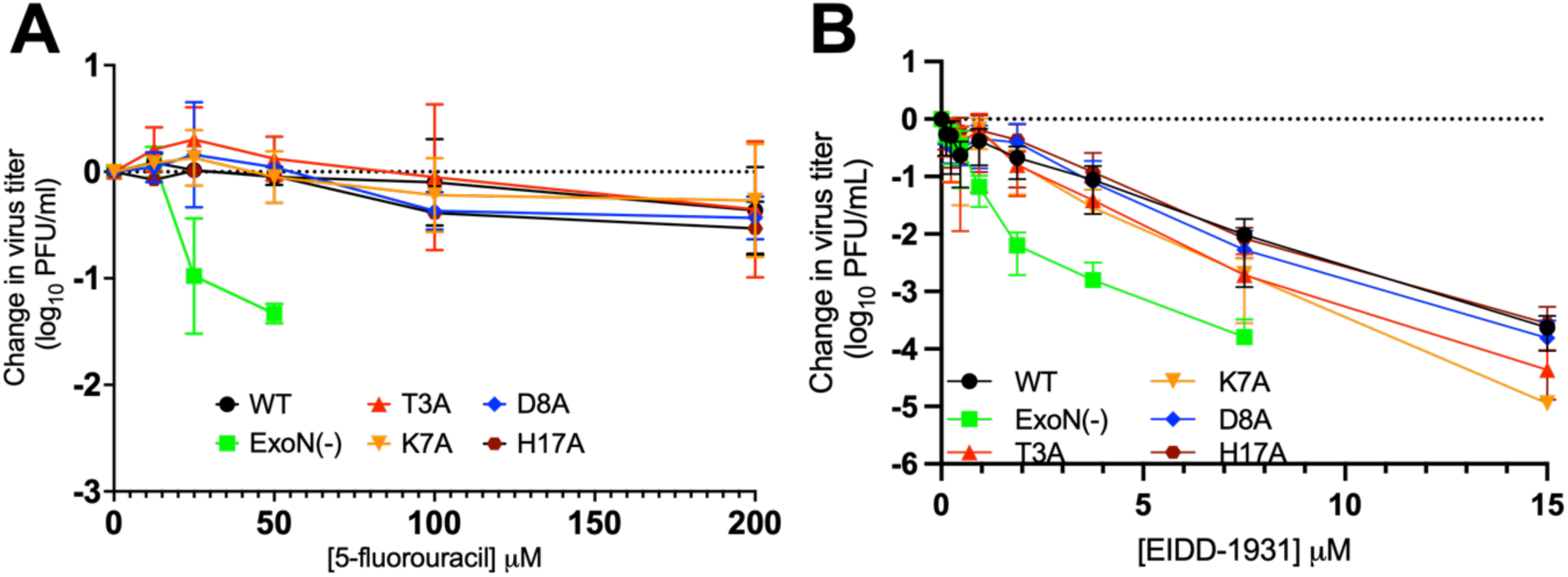
Nucleoside analog sensitivity of nsp14-10 interface mutants. **(A)** Change in viral titer in response to varying concentrations of the nucleoside analog 5-fluorouracil (5-FU). DBT-9 cells were infected with the indicated virus at an MOI of 0.01 PFU/cell and treated with the indicated concentration of 5-fluorouracil. Supernatant samples were collected 24 hpi and virus titer was determined by plaque assay. Points represent experimental means (n=2) ± SEM. **(B)** Change in viral titer in response to varying concentrations of the active form of the nucleoside analog molnupiravir, EIDD-1931, for the indicated viruses. DBT-9 cells were infected with the indicated virus at an MOI of 0.01 PFU/cell and treated with the indicated concentration of EIDD-1931. Supernatant samples were collected 24 hpi and virus titer was determined by plaque assay. Points represent experiment means (n=3) ± SEM.

### Biochemical exonuclease activity of nsp14-10 interface mutants

To test the exonuclease activity of nsp14 T3A, K7A, D8A, and H17A mutants, we performed *in vitro* exonuclease activity experiments using a recombinant MHV nsp10/14 fusion protein biochemical assay adapted from Canal et al. 2021 (39) (Fig. 5). WT and mutant MHV nsp10/14 proteins were expressed, purified, and incubated with a double-stranded RNA molecule containing a 5’-TexasRed fluorophore and 3’-quencher. Exonuclease activity was quantified by fluorescence readout over time with increasing signal indicating increased exonuclease activity. The WT MHV nsp10/14 fusion protein generated robust signal, while the MHV-ExoN(-) mutant (D89A/E91A) protein signal was indistinguishable from probe alone, confirming the complete biochemical inactivation of nsp14-ExoN. (Fig.5). The T3A, K7A, D8A, and H17A mutants each demonstrated retention or impairment of exonuclease activity, consistent with the impact of mutations on replication and fitness, but did not correlate with the retention of native resistance to nucleoside analogs 5-FU and EIDD-1931. These data indicate that nsp14 mutations at the nsp10 interface can impair nsp14 exonuclease *in vitro* activity to varying degrees, but also suggest that exonuclease impairments may be decoupled from viral sensitivity to mutagenic nucleoside analogs.

**Figure 5.**
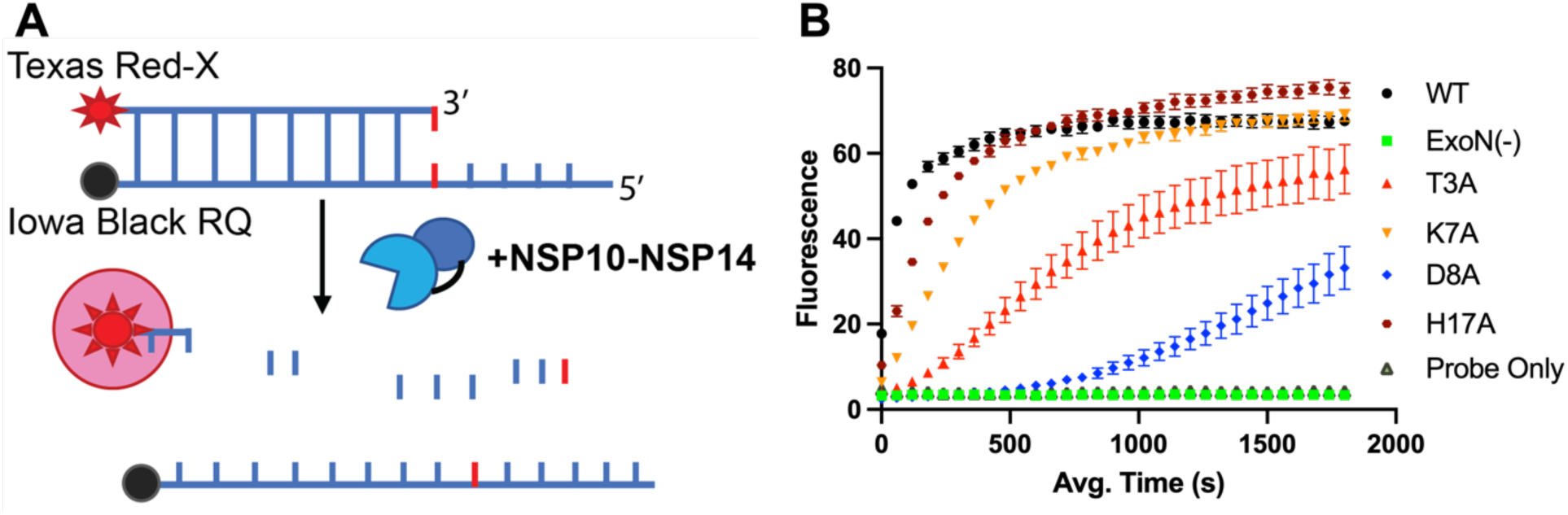
RNA cleavage of nsp14-10 interface mutants. **(A)** The *in vitro* exonuclease activity was determined by a fluorescence RNA cleavage assay using nsp10/14 fusion proteins encoding interface substitutions. The Texas red fluorescence increases with increased exonuclease activity on the partially DS RNA template. **(B)** The RNA cleavage activity of WT MHV nsp14, nsp14-ExoN(-) and interface mutants was measured over time using normalized amounts of protein and 250 nM probe. Points represent experiment means (n=4) ± SEM.

### Passage adaptation of nsp14 interface mutants K7A and D8A

Based on the significant impairment of D8A and K7A mutant replication and exonuclease activities, we passaged these viruses to identify potential intra-and intermolecular interactions of interface residues (Fig. 6). Starting with the P0 viral stocks, we serially passaged WT MHV alongside K7A and D8A mutant viruses in DBT-9 cells. Infected-cell supernatants were passaged 15 times, by which all virus populations demonstrated similar cytopathic effect (syncytia formation and cell loss) at the time of harvesting. We then compared the replication kinetics and competitive fitness of mutant passage population viruses at passage 1 (P1) and 15 (P15) to WT MHV (Fig. 6). For both the K7A and D8A mutants, passaging selected for a more rapid onset of exponential replication so that by P15, both populations demonstrated replication kinetics and peak titers closer to or indistinguishable from WT MHV. Both D8A and K7A P15 populations had higher relative fitness compared to the P1 viruses, but remained less fit compared to WT.

**Figure 6.**
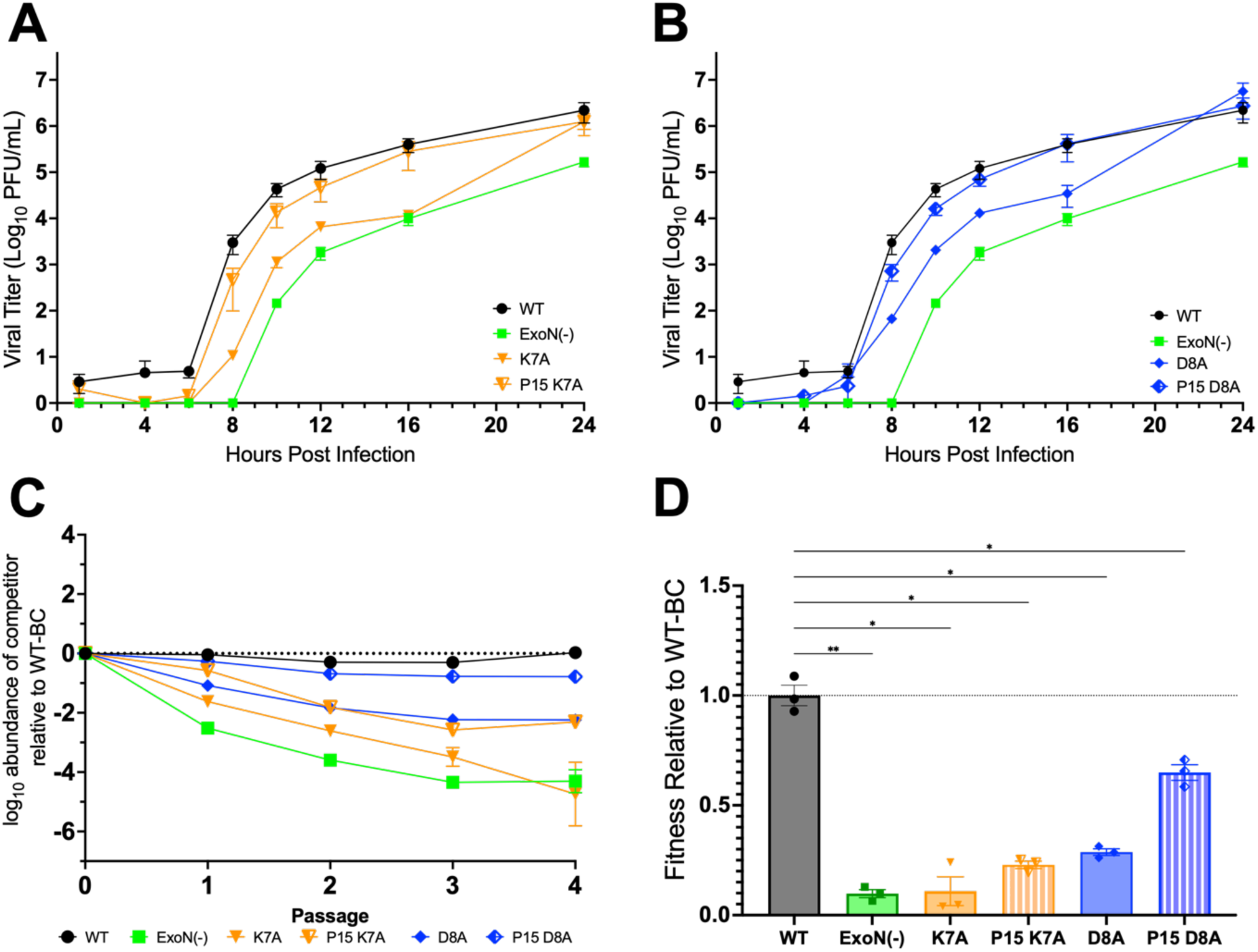
Replication and fitness of nsp14-10 interface mutant passage populations. **(A, B)** DBT-9 cells were infected with the indicated virus at an MOI of 0.01 PFU/cell. Supernatant samples were collected at the indicated times, and the virus titer was determined by plaque assay. Data shown are from three independent experiments, each with three replicates. Points represent experiment means (n=3) ± SEM. Replication of K7A P1 and P15 **(A)** and D8A P1 and P15 **(B)** compared to WT and ExoN(-). **(C)** Competitive fitness of nsp14 mutant passage population viruses. Graphed are mean values from three independent lineages. **(D)** Linear regression from panel C was used to determine the relative fitness for each nonbarcoded virus. Individual data means are graphed (n=3) ± SEM. Statistical significance determined by one-way ANOVA with Dunnett’s multiple-comparison test. * p < 0.05; ** p < 0.01.

### Sequence analysis of passage populations

We performed RNA sequencing of the P1 and P15 virus populations to identify minority variants and other mutations across the genome that arose during passaging (Table 1). Passage of WT-MHV showed low-level tissue culture adaptive changes in E and M structural proteins but no detectable coding changes with frequencies above 0.01 in any replicase nonstructural protein. The K7A mutant also had low level (<0.12) aa substitutions in E and S, and the D8A mutant had a single substitution in ORF4a at a frequency of 0.17. Both K7A and D8A retained their primary introduced changes and selected for other substitutions in nsp14. The K7A mutant passage selected an nsp14 V82I substitution (P1-0.23 and P15-0.76) as well as P15 substitutions >0.1 in nsp1, nsp3 and nsp8. The D8A mutant passage selected for three substitutions in nsp14: same-site substitution D8V (P15-0.23); T49N (P1-0.07, P15-0.41); and R52Q (P1-0.18, P15-0.12) as well as P15 substitutions >0.1 in nsp12 and nsp13. Because all three adaptive nsp14 substitutions were present at P1 in K7A and D8A, we performed Sanger sequencing of P0 recovered virus from infected cell lysates, which showed a minor peak encoding V82I along with V82 in the K7A P0. In contrast, theT49N and R52Q substitution mutations were not observed at P0. Sanger sequencing confirmed that the V82I mutation was not present in the cloned K7A cDNA fragment used to generate the virus. These results are the first to show selection of mutations in nsp14 proximate and more distant from the interface, and their early appearance suggests they may have been necessary for efficient growth of the K7A and D8A viruses during recovery. This significant selective pressure was also supported by the appearance of the same site D8V substitution which was not detected at P0 or P1 but emerged by P15.

**Table 1.**
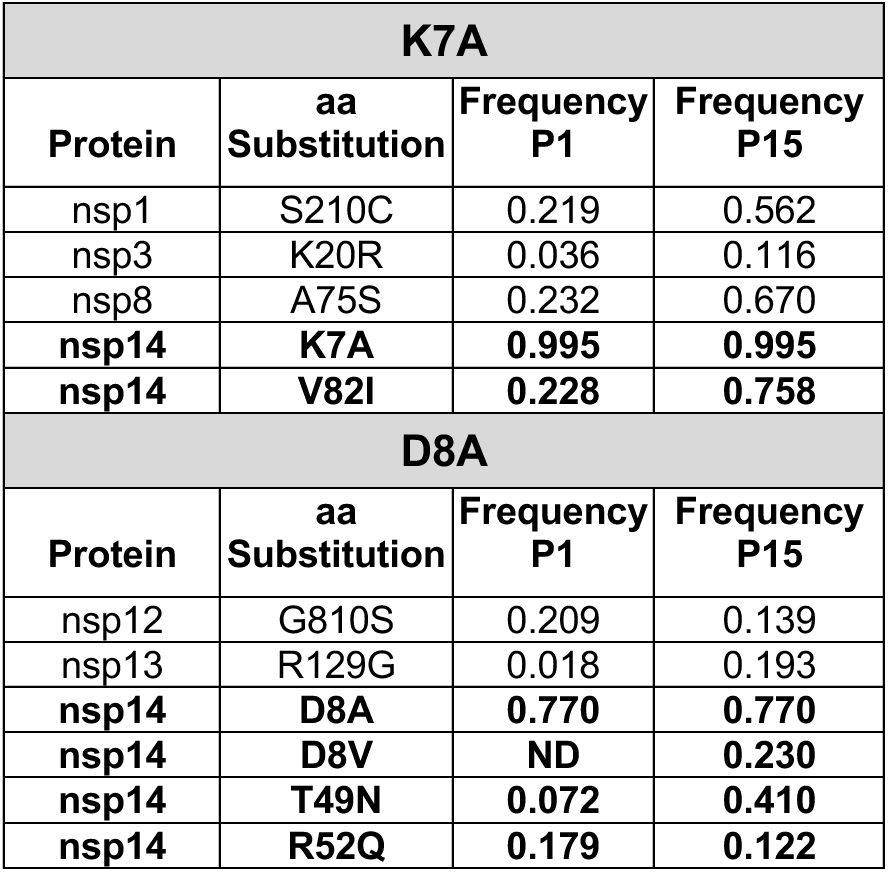
RNA Seq analysis of K7A and D8A passage populations. The nsp14 mutants were serially passaged 15 times. DBT-9 cells were infected with the indicated population virus and RNA was extracted from infected cell monolayers and sequenced. Coding mutations across the genome with >0.1 frequency are reported, showing the resultant substitution, location, and frequency.

| K7A |  |  |  |
| --- | --- | --- | --- |
| Protein | aa Substitution | Frequency P1 | Frequency P15 |
| nsp1 | S210C | 0.219 | 0.562 |
| nsp3 | K20R | 0.036 | 0.116 |
| nsp8 | A75S | 0.232 | 0.670 |
| **nsp14** | **K7A** | **0.995** | **0.995** |
| **nsp14** | **V82I** | **0.228** | **0.758** |
| D8A |  |  |  |
| Protein | aa Substitution | Frequency P1 | Frequency P15 |
| nsp12 | G810S | 0.209 | 0.139 |
| nsp13 | R129G | 0.018 | 0.193 |
| **nsp14** | **D8A** | **0.770** | **0.770** |
| **nsp14** | **D8V** | **ND** | **0.230** |
| **nsp14** | **T49N** | **0.072** | **0.410** |
| **nsp14** | **R52Q** | **0.179** | **0.122** |

### Replication of K7A and D8A with engineered nsp14 substitutions

We tested the contribution of the passage-selected nsp14 substitutions on replication by engineering the changes in the presence of K7A or D8A and recovering mutant viruses. We compared WT-MHV to the original engineered mutations (K7A and D8A), their P15 populations, and the viruses encoding K7A-V82I, D8A-T49N, D8A-R52Q, and D8A-T49N-R52Q (Fig. 7). The K7A-V82I virus had intermediate replication between K7A and the P15 K7A population. D8A-T49N, D8A-R52Q, and D8A-T49N-R52Q all had replication intermediate between D8A and D8A-P15 populations, with the exception that they achieved titers equal to D8-P15 and WT-MHV by late times of infection (16hpi). For D8A, the results show that either T49N or R52Q alone is sufficient to improve the impaired replication kinetics of D8A, but there is not an additive effect from both mutations in a D8A background. These data, paired with the frequency of T49N and R52Q, suggest that these substitutions may have been selected independently, and D8A-T49N-R52Q may not represent the genotype for individuals in the population. Overall, the experiments show that for both the K7A and D8A mutants, the selected mutations in nsp14 alone or together contribute to the adaptation of replication in isolation from other selected mutations or tissue culture adaptation.

**Figure 7.**
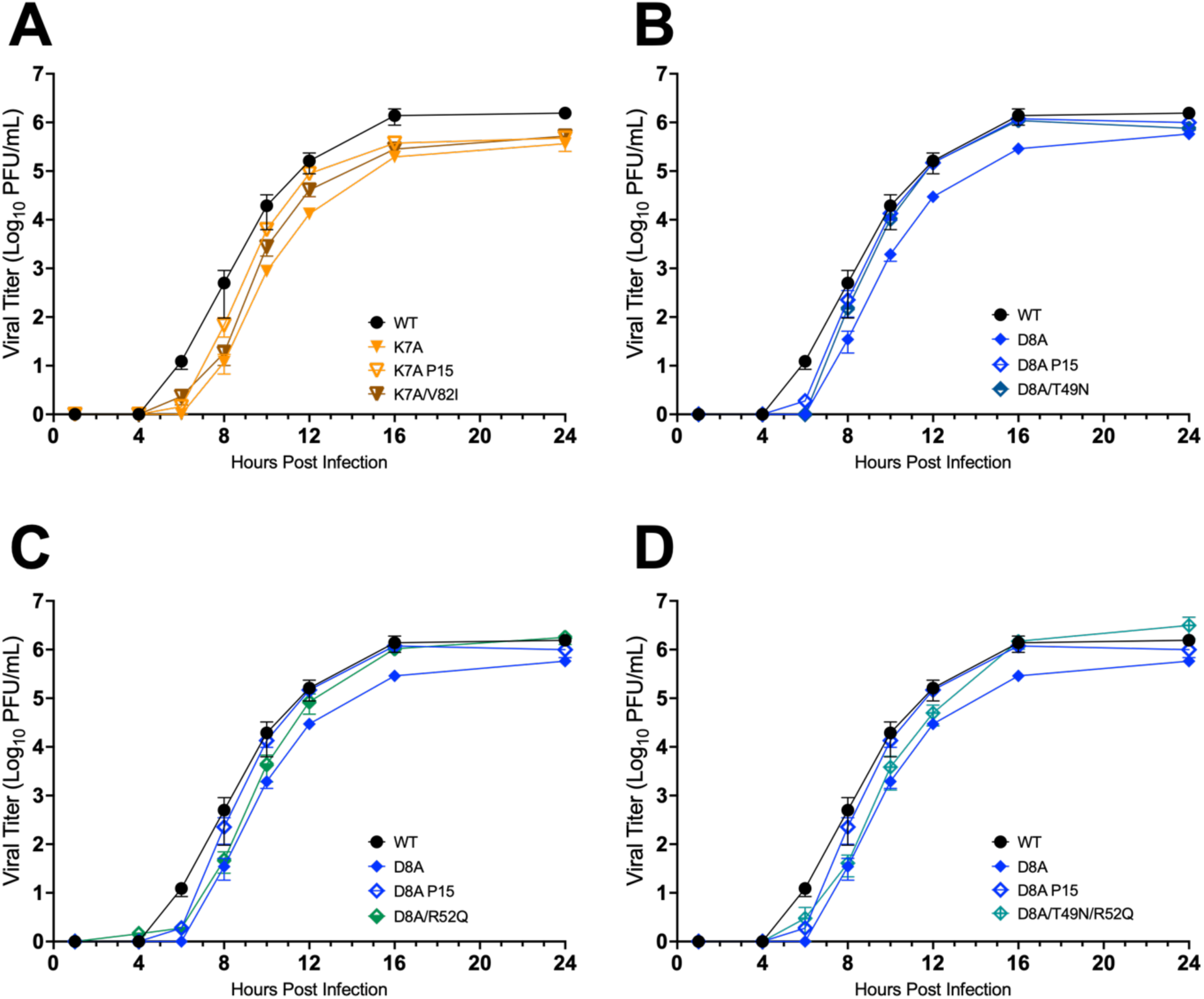
Replication of engineered nsp14-10 interface mutants. Replication of WT MHV and mutant viruses. DBT-9 cells were infected with the indicated virus at an MOI of 0.01 PFU/cell. Supernatant samples were collected at the indicated times, and the virus titer was determined by plaque assay. Data shown are from three independent experiments, each with three replicates; data is separated into four panels for ease of viewing. Points represent experiment means (n=3) ± SEM. **(A)** K7A, K7A-P15, and K7A-V82I. **(B)** D8A, D8A-P15, and D8A-T49N. **(C)** D8A, D8A-P15, and D8A-R52Q. **(D)** D8A, D8A-P15, and D8A-T49N-R52Q.

### Exonuclease activity of nsp14 adaptive mutants

Finally, we determined the impact of the nsp14 passage-adaptive mutants on the exonuclease activity of nsp14. We generated nsp10/14 fusion constructs with K7A-V82I substitutions and D8A-T49N-R52Q substitutions and compared the exonuclease activity to K7A, D8A, and WT under the same conditions listed above (Fig.8). The K7A-V82I and D8A-T49N-R52Q both had increases in total fluorescence compared to K7A and D8A, respectively, indicating the ability of these residues to partially compensate for the defects in exonuclease activity of the K7 and D8A mutants; however, neither construct approached WT-levels of exonuclease activity. These data suggest that passaging nsp14 interface mutant viruses can act as selective pressure to identify residues outside of the canonical catalytic domains that impact exonuclease activity and other functions of the nsp14 ExoN domain.

**Figure 8.**
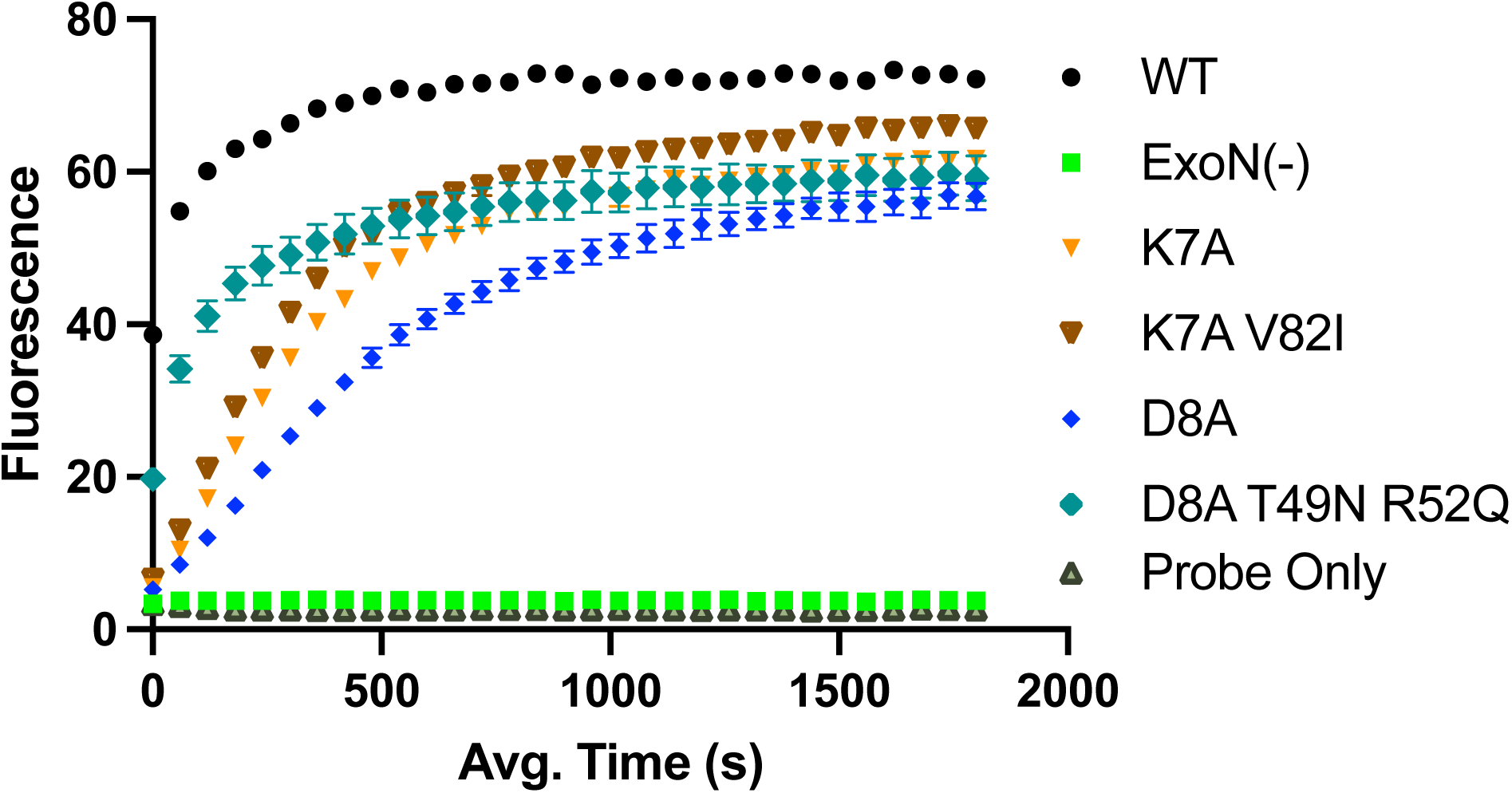
RNA cleavage of nsp14 adaptive mutants. The *in vitro* exonuclease activity was determined by a fluorescence RNA cleavage assay using 500 nM WT protein or mutant nsp10/14 fusion protein normalized to WT by SDS PAGE mixed with 250 nM of probe. Points represent experiment means (n=4) ± SEM.

## DISCUSSION

Here we show that amino acid substitutions in the MHV nsp14 at the interface with nsp10 result in varying degrees of impairment of replication, fitness, and biochemical exonuclease activity. Solved structures of nsp14/10 complexes and biochemical experiments implicate multiple contact residues between the two proteins (27–29). Our data demonstrate that individual nsp14 residue changes may not completely abolish the interactions of nsp14 and nsp10. Future experiments are needed to fully map the requirements of each interface residue, and it is possible that changes at other interface residues not tested here are absolutely required for protein-protein interaction and particle viability. For example, the same-site selection of D8V in the D8A P15 population suggests that other residues at the location may be tolerated and/or restore the defect. Nonetheless, these experiments contribute to the growing understanding of the complex protein-protein interactions responsible for coronavirus replication.

Many studies of CoV nsp14 biochemical and replication activities have focused on the exoribonuclease domain catalytic motifs and have shown nsp14 functions during RNA synthesis, genome replication fidelity, subgenomic mRNA transcription, and interferon antagonism (3,5,15,16,19,27,33,37). The well-characterized MHV nsp14 catalytic motif I mutant virus D89A/E91A (ExoN-) and homologous SARS-CoV ExoN(-) mutant have been powerful tools for probing nsp14 and CoV RTC functions. However, the inability, to date, from many labs in generating the corresponding ExoN(-) mutants in human pathogenic CoVs including MERS-CoV, SARS-CoV-2, and HCoV-229E has limited our ability to understand conserved and unique nsp14 function across divergent CoVs (14,26). Our results in this study indicate that targeting the nsp14-10 interface can yield viral and biochemical phenotypes intermediate between WT and ExoN(-). Thus, genetic alterations at the nsp14-10 interface may provide a novel approach for experiments aimed at understanding the functions of nsp14 in pathogenic CoVs.

Our passage experiments of K7A and D8A mutant viruses were designed to add selective pressures for improved replication; 15 passages were sufficient for partial restoration of replication, exonuclease activity, and fitness. The resultant D8A-P15 population harbored T49N and R52Q coding changes while the K7A-P15 population harbored a V82I coding change. Since the structure of MHV nsp14 has not been reported, we performed ColabFOLD *de novo* modeling of the MHV nsp14 based on the solved structure of the SARS-CoV-2 nsp14 (PDB: 7N0B) (Fig. 9). Structural alignment of MHV and SARS-CoV-2 nsp14 (RMSD of 1.36) indicate that T49 and R52 residues are located proximal to the D8 residue near other residues reported to interact with nsp10. In contrast, the V82 residue is located distal from the nsp10 interface near the N7-MTase domain of nsp14. The results suggest that there are both close and long-range communication nodes in the ExoN domain that determine its function and that multiple intra or intermolecular adaptive pathways are possible with nsp14/10 interface mutations, and they suggest that the CoV RTCs may adopt multiple conformations and protein-protein interactions during replication. This is supported by our recent report showing that viable mutants of MHV can be recovered with deletions of the cleavage sites at the amino and carboxy termini of nsp14, resulting in nsp13-14 and nsp14-15 obligate fusion proteins in the CoV RTC (36).

**Figure 9.**
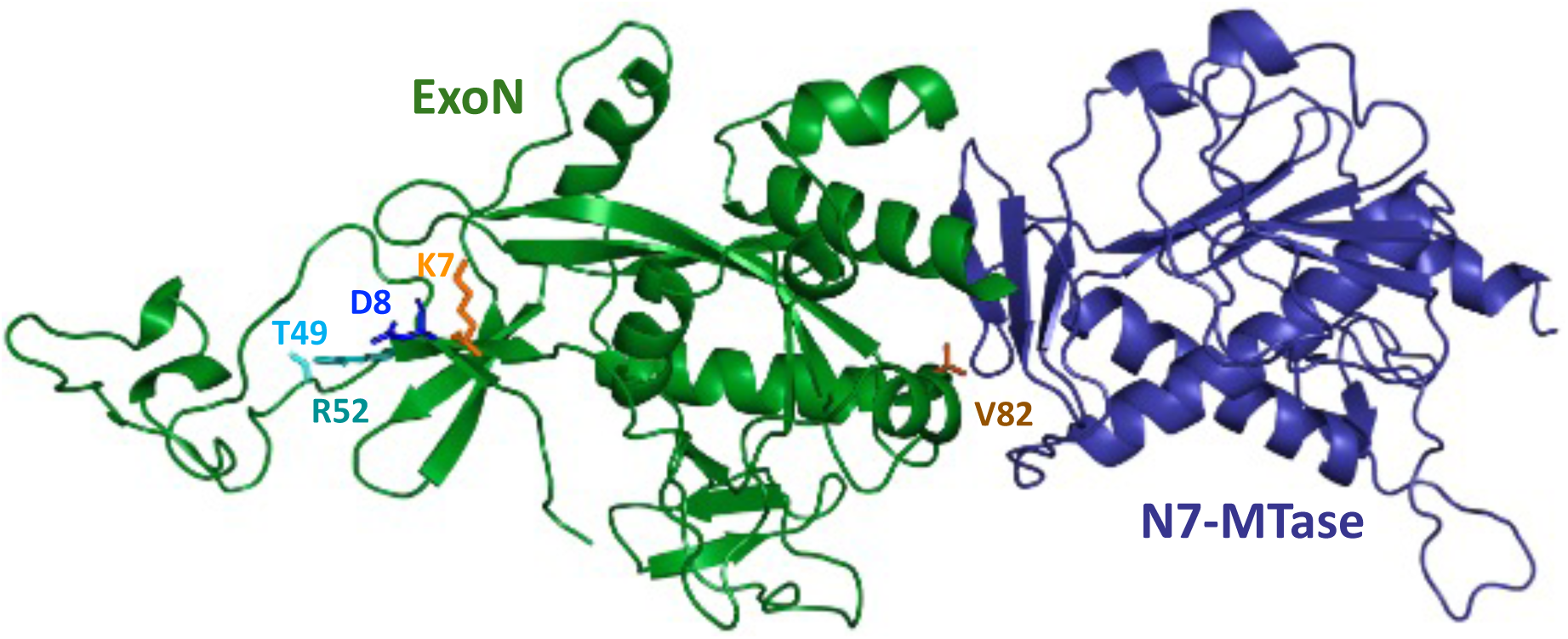
*De novo* model of MHV nsp14. PyMOL structure of the ColabFold *de novo* model of the MHV nsp14 showing the 3’-5’ exoribonuclease [ExoN] (green) and N7 methyltransferase [N7 MTase] (blue) domains. The sites of alanine mutagenesis are shown by colored side chains K7A (orange) and D8A (blue). The residues with mutations identified by Sanger sequencing and RNA sequencing are also shown: T49 and R52 residues (D8A passage) shown in cyan and teal. V82 residue (K7A passage) is shown in brown.

Our previous studies with MHV-ExoN(-) mutant viruses demonstrated a correlation between exonuclease activity and native resistance to mutagenic nucleoside analogues such as 5-FU (5). Experiments in this study demonstrate that biochemical exonuclease activity of nsp14-10 interface mutants could be significantly impaired while retaining WT-like resistance to mutagenic nucleoside analogs 5-FU and EIDD-1931. This decoupling of exonuclease activity and mutagen sensitivity was thus unexpected and raises important questions about the role of nsp14-10 interactions in the replication-transcription complex in relation to nucleotide sensitivity and proofreading. Possibilities for investigation include changes in how nsp14 interacts with the nsp12-RdRp for selectivity, or that nucleotide removal is only altered once the exonuclease activity is reduced below a threshold.

A biochemical assay using an nsp10/nsp14 fusion protein was previously developed using SARS-CoV-2 proteins, in part to help identify nsp14 inhibitors (39). The MHV nsp10/14 fusion protein system and exoribonuclease assay described here is a powerful tool which can be used as a parallel model with MHV genetic and *in vivo* studies to compare enzymatic activity with viral activity during infection in a useful BSL-2 CoV model. This fusion protein can be adapted to examine functions such as exoribonuclease activity on WT RNAs, RNA with incorporated nucleoside analogs, or in the context of nsp10 or nsp14 inhibitors. Further, this MHV nsp10/14 fusion protein may serve as the basis for a buildable *in vitro* model of the CoV RTC by incorporating multiple replication nsps or using additional approaches like magnetic tweezers for real-time analysis of the role of nsp14-10 with multiple other replicase proteins during RNA synthesis, capping, and proofreading (40). Together, these results demonstrate for the first time the contribution of nsp14 residues to the nsp14-10 interface and importance in virus replication and fitness, identify the interface as a compelling target for antiviral drugs, and introduce a novel tool for parallel genetic and biochemical analyses in a model CoV.

## MATERIALS AND METHODS

### Cell culture

Murine astrocytoma delayed brain tumor clone 9 (DBT-9) cells and baby hamster kidney cells stably expressing the MHV receptor (BHK-R) were maintained at 37°C in Dulbecco’s modified Eagle medium (DMEM; Gibco), supplemented with 10% fetal bovine serum (FBS; Invitrogen), 100 U/ml penicillin and streptomycin (Gibco), 10 mM HEPES buffer (Corning), and 0.25 μg/mL amphotericin B (Corning). BHK-R cells were also supplemented with 0.8 mg/mL G418 sulfate (Corning). Cells were routinely washed with Dulbecco’s phosphate-buffered saline without calcium chloride or magnesium chloride (PBS −/−). Cells were detached during passage and expansion with 0.05% trypsin-EDTA (Gibco).

### Viruses and amino acid conservation

All work with murine hepatitis virus (MHV) was performed using the recombinant WT strain MHV-A59 (GenBank accession number AY910861.1). MHV infectious clones were used as templates for mutagenesis and infection experiments (41). Site-directed mutagenesis by “round-the-horn” PCR was used to generate substitutions at the indicated sites (42). MHV infectious clone F fragment was used as a template to substitute nucleotides 18,167-18,169 (nsp14 T3A), 18,179-18,181 (nsp14 K7A), 18,182 -18,184 (nsp14 D8A), 18,209-18,211 (nsp14 H17A), 18,305 -18,307 (nsp14 T49N), 18,314-18,316 (nsp14 R52Q), 18,404-18,406 (nsp14 V82I), using the following primers: T3AF (5’-GCCAATTTGTTTAAGGATTGTAGCAGGAG-3’), T3AR (5’-AGTACACTGTAATCGTGGATTGTTAAT-3’); K7AF (5’-GCCGATTGTAGCAGGAGCTATGTAG-3’), K7AR (5’-AAACAAATTTGTAGTACACTGTAATCGT-3’); D8AF (5’-GCATGTAGCAGGAGCTATGTAGG-3’), D8AR (5’-CTTAAACAAATTTGTAGTACACTGTAATCGT); H17AF (5’-GCACCAGCCCATGCAC-3’), H17AR (5’-ATATCCTACATAGCTCCTGC-3’); T49NF (5’-CTATTCGCGGCTTATATCACTC-3’), T49NR (5’-TTGACAGCAGAATCAGCAAC-3’); R52QF (5’-ACTTATATCACTCATGGGATTCAAGC-3’), R52QR (5’-TGCGAATAAGTGACAGCAGAATCAG-3’); V82IF (5’-ACAGAGCCTGGGTTGG-3’), V82IR (5’-TACGTTTGATAGCTTCATCTCT-3’); K7A/D8AF (5’-GCCGCATGTAGCAGGAGCTATG-3’), K7A/D8AR (5’-AAACAAATTTGTAGTACACTGTAATCGTG-3’). All primers were 5’-phosphorylated with T4 polynucleotide kinase using an ATP-containing reaction buffer (NEB). Template backbone DNA was digested with DpnI (NEB), and amplified DNA was separated by electrophoresis and extracted from agarose (Promega). Ligated DNA was transformed into Top10 competent *Escherichia coli* cells (Thermo) and amplified in liquid culture, and sequences were confirmed by Sanger sequencing. Assembly and recovery of recombinant MHV has been described previously (41). Electroporated cells were monitored for cytopathic effect (CPE) and cell flasks were frozen at -80°C when approximately 80% of the monolayer was involved in CPE. Cells were thawed, debris was pelleted, and virus-containing supernatants were aliquoted and stored at -80°C (Passage 0). The P0 stocks were used for all experiments above, except for T3A virus, in which case the P1 stocks were used. Engineered mutations of the P0 or P1 stocks used for experiments were confirmed by Sanger sequencing. Samples of viral stock supernatants were collected in TRIzol (Ambion), and viral RNA was extracted by chloroform extraction and purified using the KingFisher MagMAX Viral/Pathogen Nucleic Acid Isolation Kit (Thermo). Viral cDNA was generated with SuperScript IV reverse transcriptase (Thermo) using random hexamers and oligo(dTs). Amplicons (each 3-4 kbs in length) were generated via PCR using EasyA polymerase (Agilent) and Sanger sequenced.

### Amino acid sequence conservation

To determine conservation across multiple coronaviruses, multiple sequence alignments were generated using MacVector. The following sequences were used for reference to compare amino acid identity by NCBI Basic Local Alignment Search Tool (BLAST): MHV_A549 (YP_ 009915697.1), SARS-CoV-2 (YP-009725306.1).

### Virus replication assays

DBT-9 cells were plated at a density of 6e5 cells per well 24 hours before infection. Cells were then infected at an MOI of 0.01 PFU/cell for 1 hour. Inocula were removed, and the cells were washed with PBS before addition of prewarmed medium. Supernatants were harvested at the indicated times post-infection and titers were determined by plaque assay.

### Plaque Assays

Plaque assays were performed in sub-confluent DBT cells seeded in 6-well plates. Serial dilutions were plated in duplicate and overlaid with 1% agar in DMEM. Titers were scored at 24 hpi.

### Competitive fitness assay

The MHV competitive fitness assay was previously described in detail (36–38). Briefly, sub-confluent DBT-9 cells were co-infected with the indicated virus and a barcoded WT-MHV reference virus with seven silent mutations in nsp2 (1301-CAGCAGT-1307) at a total MOI of 0.1 PFU/cell (0.05 MOI for each virus) in three independent lineages. The resulting virus was passaged four additional times, each at a constant MOI of 0.1 PFU/cell. Viral RNA from each passage supernatant was extracted in TRIzol and purified with a KingFisher II (ThermoFisher Scientific) according to the manufacturer’s protocol. RNA corresponding to the barcoded WT reference and test viruses was determined by one-step RT-qPCR using SYBR green. BC WT reference RNA was detected with forward (5′-*CTATGCTGTATACGGACAGCAGT*) and reverse (5′-*GGTGTCACCACAACAATCCAC*) primers, and test virus RNA was detected with forward (5′-*CTATGCTGTATACGGATTCGTCC*-3′) reverse (5′-*GGTGTCACCACAACAATCCAC*) primers using a Power SYBR green RNA-to-Ct 1-step kit (Applied Biosystems) on a StepOnePlus real-time PCR system (Applied Biosystems). The log-transformed cycle threshold (*C_T_*) ratio of test versus reference was plotted over passage, and relative fitness was determined by comparing the slopes of linear regression.

### Nucleoside analogue sensitivity studies

For 5-fluorouracil sensitivity assays, DBT-9 cells were pre-incubated with the indicated concentration of 5-fluorouracil or DMSO control for 30 minutes. Sub-confluent monolayers of DBT-9 cells were infected with MHV at an MOI of 0.01 PFU per cell for 1 h at 37°C. The inoculum was removed and replaced with medium containing the indicated compound concentration (5-fluorouracil (Sigma) or EIDD-1931 (MedChemExpress)). Cell supernatants were harvested 24 h post-infection. Titers were determined by plaque assay as previously described.

### Passaging nsp14 mutant viruses

WT MHV, MHV with the nsp14 K7A substitution, and MHV with the nsp14 D8A substitution were each passaged in T25 flasks, and infection was initiated 16 hours after plating using 1 mL of passage 0 stock virus. Virus supernatants were harvested 8 hpi and stored on ice overnight at 4°C, and total RNA from virus-infected monolayers was harvested using TRIzol (Invitrogen). A constant volume of 1 mL was used to initiate subsequent passages for a total of 15 passages.

### Illumina RNA-sequencing of viral RNA, processing and alignment

Total RNA was extracted from monolayers infected with WT and mutant MHV viruses at passage 1 (P1) and passage 15 (P15) using TRIzol (Invitrogen) according to the manufacturer’s instructions. For RNA-Seq, total RNA underwent poly(A) selection followed by NovaSeq PE150 sequencing (Illumina) at 15 million reads per sample at the Vanderbilt University Medical Center (VUMC) core facility, Vanderbilt Technologies for Advanced Genomics (VANTAGE). VANTAGE performed base-calling and read demultiplexing. The CoVariant pipeline (43) was used for variant analysis. The first module trims and aligns raw FASTQ files to the viral genome for each specified sample using a standard Bash shell script. To summarize, raw reads were processed by first removing the Illumina TruSeq adapter using Trimmomatic. Reads shorter than 36 bp were removed, and low-quality bases (Q score of <30) were trimmed from read ends. The raw FASTQ files were aligned to the MHV-A59 genome (AY910861.1) by using the CoVariant Python3 script command line parameters. For variant analysis, the sequence alignment map (SAM) file was processed using the samtools suite and alignment statistics output was generated by samtools idxstats to an output text file. Nucleotide depth at each position was calculated from the SAM files using BBMap (Bushnell) pileup.sh.

### NSP10/14 fusion cloning protein expression

A MHV (NP_045299) nsp10/14 fusion containing a 2x GGS linker between nsp10 and nsp14 was codon optimized for expression in *E. coli* using IDT codon optimization tool and cloned into a pK27 vector that contains an N-terminal 6x-His-tag, Flag-tag, and a SUMO-tag as originally described for SARS-CoV-2 (39). Plasmids were transformed into C41 (DE3) pLysS competent cells (Sigma-Aldrich CMC0018). Cultures were grown in terrific broth with 50 µg/mL kanamycin at 37°C until they reached an OD600 between 0.8 and 1, and then cooled at 4°C for one hour. Cultures were induced with 0.5 mM IPTG and incubated overnight at 18*°*C. The following day, cells were centrifuged, resuspended in 25 mL of lysis buffer (50 mM NaH_2_PO_4_, 300 mM NaCl, 10 mM imidazole, pH 8, complete EDTA-free protease inhibitor tablet from Sigma Aldrich (04693132001)), lysed with a sonicator, recentrifuged at 18,000 x g for 30 minutes at 4°C, and the supernatant was filtered through a 0.45 um filter. After filtering, supernatant was resuspended 1:5 with lysis buffer and incubated at 4°C with Takara’s Talon affinity resin. Resin was then collected via gravity chromatography and eluted with elution buffer (50 mM NaH_2_PO_4_, 300 mN NaCl, 500 mM imidazole pH 8). The protein was then dialyzed twice against PBS to remove traces of imidazole and concentrated in a 50K molecular weight PES concentrator (Thermo Scientific, 88541). Protein quantitation was carried out using Pierce Rapid Gold BCA Protein assay (Thermo Scientific, A53225) and a 4-20% mini-Protean TGX Stain-Free gel (Bio-Rad) was run to confirm protein concentrations. Gel was imaged using ChemiDoc MP Imaging system using stain free settings.

### FRET based In vitro exonuclease assay

Exonuclease activity was quantified using purified WT and mutant nsp10/14 protein diluted in NSP buffer (50mM Tris HCl, 0.0001% Tween 20, 5% glycerol, 1.5mM MgCl_2_, 20mM NaCl, 0.5mM TCEP, 0.1ug/ml BSA). 500nM WT protein was mixed with 250nM of annealed RNA oligos containing fluor and quencher 5’TexRd-XN/rArCrArArArArCrGrGrCrCrCrA and rA*rA*rA*rU*rA*rG*rG*rG*rC*rC*rG*rU*rU*rU*rU*rG*rU*/3’IAbRQSp/ (*indicates phosphothioate bond). Fluorescence was measured using a Thermo Scientific Varioskan Lux 3020-81011 plate reader iii using (excitation: 590nm, emission 615nm) for 30 minutes. For interface and adaptive mutants, protein was normalized to WT concentrations using a combination of BCA assay results and intensity of protein band imaged in a mini-Protean TGX Stain-Free gel.

### Model building of MHV nsp14

A *de novo* model of the MHV nsp14 was generated by ColabFold using the MHV_A59 sequence (YP_009915687.1). The results were then visualized and aligned to the SARS-CoV-2 nsp14 from the solved structure 7N0B in PyMOL (Schrödinger).

### Statistics

Statistical tests were performed using GraphPad (La Jolla, CA) Prism 10 software as described in the respective figure legends.

## Data Availability Statement

FASTQ files for the RNA-seq variant analysis have been deposited in the National Center for Biotechnology Information Sequence Read Archive (NCBISRA) under the accession number PRJNA1151693.

## Acknowledgements

This work was supported by NIH-NIAID grants AI108197 (MRD) and U19AI171292 (MRD and NSH).

